# ATAC and histone H3K9me3 landscapes revealed the altered epigenome by fetal-neonatal iron deficiency in the adult male rat hippocampus

**DOI:** 10.1101/2022.06.07.495122

**Authors:** Shirelle X. Liu, Aarthi Ramakrishnan, Li Shen, Jonathan C. Gewirtz, Michael K. Georgieff, Phu V. Tran

**Author notes:** Corresponding Author: Phu Tran, PhD, 420 Delaware St, SE, MMC 391 Mayo, Minneapolis, MN 55455.

## Abstract

Iron deficiency during the fetal-neonatal period results in long-term neurodevelopmental impairments associated with pervasive and widespread hippocampal gene dysregulation. Globally, fetal-neonatal iron deficiency produces both long-term activation and repression of hundreds of loci in the adult rat hippocampus. Prenatal choline (a methyl donor) supplementation can partially reverse these effects, suggesting an interaction between iron and choline in regulating the hippocampal transcriptome. To gain insights into the underlying epigenetic signatures, we integrate hippocampal transcriptomes and epigenetic marks of active (transposase accessible chromatin/ATAC) and repressed (H3K9me3 enrichment) genes in adult rats that had been exposed to fetal-neonatal iron deficiency with or without prenatal choline supplementation. Rats were made iron-deficient during fetal and neonatal period by limiting maternal iron intake from gestational day (G) 2 through postnatal day (P) 7. Choline (5.5 g/kg) was given to half of the pregnant dams during G11-18. This paradigm produced four comparison groups (Iron-sufficient [IS], Iron-deficient [ID], IS+choline [ISch], and ID+choline [IDch]). Hippocampi were collected from P65 males and analyzed for changes in chromatin conformation and histone H3K9me3 enrichment. ATAC-seq results accounted for 22% and 24%, whereas H3K9me3 enrichment accounted for 1.7% and 13% of differences in ID- and IDch-altered gene expression. These epigenetic changes were annotated onto gene networks regulating synaptic structure and plasticity, neuroinflammation, and reward circuits. The low correlation between gene dysregulation and changes in ATAC or H3K9me3 signatures indicate involvements of other epigenetic modifications. This study provides a genome-wide findings of stable epigenetic changes and lays a foundation for further analyses to elucidate more fully iron-dependent epigenetic mechanisms that underlie iron deficiency, choline supplementation, and their interactions in mediating long-term neural gene dysregulation.

**SIGNIFICANCE STATEMENT:** Early-life iron deficiency can lead to long-term neurocognitive dysfunction and persistent neural gene dysregulation, despite prompt iron replenishment, suggesting that iron deficiency results in long-term neuroepigenomic changes. This study combined RNA-seq, ATAC-seq, and ChIP-seq to provide the epigenetic basis for gene dysregulation due to fetal-neonatal iron deficiency and prenatal choline supplementation. We found that early-life iron deficiency alters epigenetic regulation of genes involved in neuronal development, cell signaling, neuroinflammation, and reward-related cognition. While choline supplementation to iron-deficient animals partially reverses these effects, it also leads to dysregulation of genes in iron-sufficient animals. The patterns of gene dysregulation were positively correlated with differences in chromatin accessibility and negatively correlated with repressive histone H3K9me3 modification. Our results indicate that these changes at the epigenetic level partially account for the long-term hippocampal gene dysregulation.

## Introduction

Iron deficiency during gestational and neonatal periods, when the brain undergoes robust growth (Kuzawa, 1998; Li et al., 2004; Fukumitsu et al., 2016), can result in short- and long-term neurodevelopmental impairments. Extensive longitudinal studies of human cohorts uncover a pattern of persistent cognitive impairment and emotional dysregulation following early postnatal iron deficiency (Lozoff et al., 2000, 2013). Animal models of fetal-neonatal iron deficiency exhibit analogous neurodevelopmental impairments that are associated with pervasive and widespread gene dysregulation in the adult hippocampus (Fretham et al., 2011; Schachtschneider et al., 2016; Tran et al., 2016; Barks et al., 2018, 2019), implicating stable changes in the epigenome.

Choline is an essential nutrient for brain development and can act as a methyl donor to modify DNA methylation and histone methylation (Cooney et al., 2002; Fisher et al., 2002; Zeisel, 2004; Davison et al., 2009; Krzysztof Blusztajn and J. Mellott, 2012; Jirtle, 2014). Choline supplementation can be beneficial in mitigating the deleterious developmental effects of early life adverse exposure and aberrant genetic factors on brain development (Wozniak et al., 2013; Mellott et al., 2017; Chin et al., 2019; Agam et al., 2020). In the context of fetal-neonatal iron deficiency, choline supplementation can influence hippocampal gene expression and epigenomic changes in an iron status-dependent manner (Kennedy et al., 2014, 2018; Tran et al., 2016; Liu et al., 2021). These findings suggest an interaction between these two essential nutrients in influencing the brain’s epigenetic landscape. However, genome-wide analyses of chromatin and histone modifications due to early-life iron deficiency or choline supplementation have not yet been performed.

In the present study, we employed Next-Generation Sequencing to assess global changes in accessible chromatin by Assay for Transposase Accessible Chromatin with sequencing (ATAC-seq) and gene silencing epigenetic signatures by histone H3K9me3 Chromatin Immunoprecipitation with sequencing (ChIP-seq) caused by fetal-neonatal iron deficiency, prenatal choline supplementation, and their interaction. We hypothesized that changes in accessible chromatin and H3K9me3 profiles can account for the lasting gene dysregulation in formerly iron-deficient and prenatal choline-supplemented rats and that these changes occur in gene networks regulating neurocognition and emotional behaviors.

## MATERIALS AND METHODS

### Animals

Gestational day (G) 2 pregnant Sprague-Dawley rats were purchased from Charles River Laboratories (Wilmington, MA). Rats were maintained in a 12-hr:12-hr light/dark cycle with *ad lib* food and water. Fetal-neonatal iron deficiency was induced by dietary manipulation as previously described (Tran et al., 2012; Kennedy et al., 2018). In brief, pregnant rats were given a purified iron-deficient diet (4 mg Fe/kg, TD 80396, Harlan Teklad, Madison, WI) from G2 to post-natal day (P) 7, then a purified iron-sufficient diet (200 mg Fe/kg, TD 09256, Harlan Teklad) thereafter. Both diets were similar in all respects except for their iron (ferric citrate) content. Control iron-sufficient rats were generated from pregnant rats maintained on an iron-sufficient diet. Half of the dams on iron-sufficient or iron-deficient diet received dietary choline supplementation (5.0 g/kg choline biartrate (Glenn et al., 2008; Wong-Goodrich et al., 2008), Iron-sufficient + choline: TD 1448261, Iron-deficient + choline: TD 110139) from G11-18, while the remaining dams received iron-modified diets with standard choline content (1.1 g/kg).

Thus, dams and their litters were randomly assigned to the following groups: iron-deficient with choline supplementation (IDch), iron-deficient without choline supplementation (ID), always iron-sufficient with choline supplementation (ISch), and always iron-sufficient without choline supplementation (IS). All litters were culled to 8 pups at birth. Only male offspring were used in experiments. To avoid litter-specific effects, two male rats per litter and ≥ 3 litters/group were used in the experiments. Rats were weaned on P21 and maintained on the iron-sufficient diet thereafter. The University of Minnesota Institutional Animal Care and Use Committee approved all experiments in this study (Protocol # 2001-37802A).

### Hippocampal dissection

P65 rats were euthanized by injection of pentobarbital (100 mg/kg, intraperitoneal). Brains were removed and bisected along the midline on an ice-cold metal block. Hippocampi were dissected bilaterally, immediately flash-frozen in liquid nitrogen, and stored at -80°C.

### Nuclei isolation and DNA transposition

Nuclei isolation and DNA transposition were carried out as previously described (Corces et al., 2017) with modifications. Briefly, flash-frozen tissue was thawed in cold homogenization buffer (5 mM CaCl_2_, 3 mM Mg(AC)_2_, 10 mM Tris pH 7.8, 0.0167 mM PMSF, 0.167 mM β-mercaptoethanol, 1x proteinase inhibitor (cOmplete), 320 mM sucrose, 0.1 mM EDTA, 0.1% CA630), and homogenized using a Pellet Pestle motor and syringe. Equal volume of 50% iodixanol solution (5 mM CaCl_2_, 3 mM Mg(AC)_2_, 10 mM Tris pH 7.8, 0.017 mM PMSF, 0.17 mM β-mercaptoethanol, 1x proteinase inhibitor (cOmplete), 50% iodixanol) was added to homogenate to reach a final concentration of 25% iodixanol. 29% (5 mM CaCl_2_, 3 mM Mg(AC)_2_, 10 mM Tris pH 7.8, 0.017 mM PMSF, 0.17 mM β-mercaptoethanol, 1x proteinase inhibitor (cOmplete), 160 mM sucrose, 29% iodixanol) and 35% (5 mM CaCl_2_, 3 mM Mg(AC)_2_, 10 mM Tris pH 7.8, 0.017 mM PMSF, 0.17 mM β-mercaptoethanol, 1x proteinase inhibitor (cOmplete), 160 mM sucrose, 35% iodixanol) iodixanol solution were sequentially added under the 25% iodixanol solution layer. The 3-layer system was centrifuged in a swinging bucket centrifuge at 4255 x g for 20 min. After centrifugation, nuclei were isolated and collected from the 29%-35% iodixanol solution interface. Isolated nuclei were counted with trypan blue (0.4%) and 50,000 nuclei were transferred into a tube containing 1 ml ATAC-RSB (10 mM Tris pH 7.8, 10 mM NaCl, 3 mM MgCl_2_) with 0.1% Tween-20. Nuclei were pelleted by centrifugation at 500 x g for 10 min. All steps above were conducted on ice or at 4°C. Nuclei were resuspended in transposition mix (25 µL 2x TD buffer (Illumina), 16.5 µL PBS, 0.5 µL 10% Tween-20, 1% digitonin, 2.5 µL Tn5 transposase (Illumina), 5 µL H_2_O), and incubated at 37°C for 30 min in a thermomixer at 1000 rpm. Transposed DNA was purified using MinElute Reaction Cleanup kit (Qiagen).

### Chromatin immunoprecipitation (ChIP)

ChIP experiments were performed as previously described (Tran et al., 2015) with modifications. In brief, chromatin was prepared from hippocampal tissue following the manufacturer’s recommendations (Millipore, Temecula, CA). Tissue was homogenized in ice-cold PBS (500 µL) using a Pellet Pestle motor and pelleted by centrifugation at 13K rpm (30s). The pellets were resuspended in PBS and cross-linked in 1% formaldehyde solution (Sigma). Following the removal of fixative and PBS rinses, lysates were resuspended in 500 µL lysis buffer (1% SDS, 10 mM EDTA, 50 mM Tris pH 8.1, 1 mM PMSF, 10 µL 10X protease inhibitor cocktails (Roche, Indianapolis, IN)), incubated in an ice bath for 10 min, and sonicated (Bioruptor Pico, Diagenode) to shear DNA. Sonicated lysates were diluted 10-fold with ChIP dilution buffer (0.01% SDS, 1.1% Triton X-100, 1.2 mM EDTA, 16.7 mM Tris-pH 8.1, 167 mM NaCl) and pre-cleared with 75 µL of Protein A agarose (50% slurry, Sigma). Pre-cleared chromatin lysate was immunoprecipitated by a ChIP-grade rabbit polyclonal antibody against histone H3K9me3 (Cat. # C15410193, Diagenode) with end-over-end rotation (4°C, overnight). The antibody-histone complex was collected through the addition of 75 µL Protein A agarose slurry with mixing (4°C, ≥ 1 hour). Following washes (per manufacturer’s protocol, Millipore), the immune-histone complex was eluted in 500 µL of elution buffer (1% SDS, 0.1 M NaHCO_3_). Reverse cross-linking was achieved by incubation in NaCl (0.2 M, 65°C, overnight). Protease digestion (20 µg proteinase K, 20 mM EDTA, 100 mM Tris-pH 6.5, 45°C, 1 hour) was performed to recover DNA, which was further purified using phenol/chloroform extraction and EtOH precipitation (1/10 volume 3 M sodium acetate, pH 5.2, 2 volume ethanol). Levels of enriched *Gapdh* (active) and *Myod1* (inactive) loci were used to validate ChIP experiments by Real-time PCR.

### Real-time PCR

For analysis of precipitated DNA from ChIP experiments, SYBR-green PCR (Fast SYBR green master mix, ABI) was used to amplify *Gapdh and Myod1*. Input DNA (10%) was used as a normalizer to account for input amount (ΔCt). Real-time PCR was performed with QuantStudio™ 5 (Thermo Fisher).

### Next-Generation Sequencing

Recovered DNA from ATAC or ChIP was delivered to the University of Minnesota Genomic Center for quality control, library preparation and sequencing. DNA was first quantified using the PicoGreen dsDNA Assay Kit (Invitrogen). ATAC-seq library preparation was performed in accordance with the ATAC-seq protocol as described by Buenrostro et al. (2015). Library preparation for ChIP-seq was completed using ThruPLEX® DNA-Seq (Rubicon Genomics) kit. Libraries were assessed for quality using electrophoresis on an Agilent Bioanalyzer (Agilent) and size-selected for 200∼800 bp fragments. Selected libraries were quantified again, and sequencing was performed at a depth of ∼46 million reads per sample using NovaSeq 6000 for ATAC-seq, and a depth of ∼25 million reads per sample using HiSeq 2500 for ChIP-seq to generate 50-bp pair-end reads.

### Bioinformatics

For ChIP-seq and ATAC-seq datasets, comprising 16 samples with 4 biological replicates in each condition (IS, ISch, ID, and IDch) were aligned to the rat reference genome (rn6) using HISAT2 (Kim et al., 2019). MACS was utilized to assess sites of chromatin accessibility in ATAC-seq samples and sites of H3K9me3 histone modification (peaks) for each of the 4 conditions (Zhang et al., 2008). diffReps (Shen et al., 2013) was run on the processed samples to find differential sites for the comparisons to the IS control group (ID, IDch, or ISch vs IS). Sequence alignments and analyses of specific loci were performed using IGV tools (Broad Institute). Changes in aligned peaks between the experimental groups (ID, IDch or ISch) and the control IS group were determined by subtraction of the aligned tracks. TFBIND (https://tfbind.hgc.jp) was used to identify potential transcription factor (TF) binding sites within a selected DNA sequence (Tsunoda and Takagi, 1999). Global analysis of overlapping between ATAC and H3K9me3 peaks was performed using 1 bp overlapping threshold to identify regions modified by both markers. For RNA-seq data, duplicate datasets from a previous study (Tran et al., 2016) were integrated and reprocessed using FastQC, Hisat2 (v2.1.0). Gene quantification was done via Feature Counts for read counts. Differentially expressed genes were identified using the edgeR (negative binomial) feature in CLCGWB (Qiagen, Redwood City, CA) and raw read counts. Each experimental group (ID, IDch or ISch) was compared to the IS control group.

### Ingenuity Pathway Analysis (IPA)

Differentially modified loci from pairwise comparisons between treatment and IS control group (e.g., ID vs IS, IDch vs IS, and ISch vs IS) were analyzed by IPA (Qiagen), a knowledge-based database of > 7.8 million findings, to identify pertinent pathways, gene networks, and biofunctions. Data were analyzed using core analyses with *p*-value ≤ 0.05. Comparisons between datasets were performed with a built-in analysis that employs the Fisher’s exact test to calculate significant effects, which is set at absolute Z-score ≥ 2.0. Positive and negative Z-scores indicate increased and decreased activity, respectively.

## RESULTS

### Iron deficiency and prenatal choline supplementation alter long-term chromatin accessibility in adult rat hippocampus

To elucidate the long-term changes in accessible chromatin structures due to fetal-neonatal iron deficiency or prenatal choline supplementation, P65 adult rat hippocampi were analyzed by ATAC-seq. Compared to the IS control group, the ID, IDch or ISch groups showed a greater number of increased than decreased accessible loci (Table 1, ATAC). Prenatal choline supplementation resulted in a higher number of accessible loci in the IDch compared to the untreated ID group (Table 1, IDch > ID). These changes occurred across the whole genome with greater proportions within the gene body and intergenic regions (Supplemental Figure 1A).

**Table 1:** Results of NGS datasets showing differentially-regulated genes between treatment groups. For ATAC-seq and ChIP-seq data, numbers of differential sites include gene body, promoter regions (proximal, +1K, +3K), and intergenic regions and numbers of loci include gene body and promoter regions that were annotated to specific genes. Selection criteria were absolute log2(Fold Change) > 0.2 and p <0.05 for RNA-seq, and false discovery rate q-value < 0.05 for ATAC-seq and ChIP-seq; n=4/group.

| NGS Data | Comparison | # Differential Sites | # Loci – Lower Enrichment | # Loci – Higher Enrichment |
| --- | --- | --- | --- | --- |
| RNA | ID vs IS | 731 | 232 | 499 |
|  | IDch vs IS | 983 | 585 | 398 |
|  | ISch vs IS | 1,102 | 489 | 613 |
| ATAC | ID vs IS | 6,001 | 1,084 | 1,586 |
|  | IDch vs IS | 7,688 | 1,274 | 2,391 |
|  | ISch vs IS | 8,789 | 1,896 | 1,973 |
| H3K9me3 | ID vs IS | 1,439 | 89 | 110 |
|  | IDch vs IS | 23,077 | 1,258 | 1,193 |
|  | ISch vs IS | 23,842 | 1,143 | 2,292 |

To determine the biological significance of the changes in accessible chromatin structures among the treatment groups, we employed IPA to functionally annotate the altered loci. Hierarchical clustering of annotated biofunctions indicated that iron deficiency increased accessibility of genes regulating quantity of cells (z-score > 2.0, Fig. 1A, ID/IS) accompanied by decreased accessibility of loci regulating neuronal maturation (z-score < -2.0, Fig. 1A, ID/IS). These changes were mitigated by choline treatment (Fig. 1A, IDch/IS). Choline treatment also decreased accessibility of genes regulating hippocampal cell death, long-term synaptic depression (LTD) of hippocampus, and gliosis of central nervous system (z-score < -2.0, Fig. 1A, IDch/IS) and increased accessibility of loci implicated in myelination, release of vesicles, and transport of mitochondria in the IDch group (z-score >2.0, Fig. 1A, IDch/IS). Choline treatment of the IS group decreased accessibility of genes regulating hippocampal formation, cell migration, synaptic structures and functions, and neuron proliferation and development (z-score < -2.0, Fig. 1A, ISch/IS).

**Figure 1:**
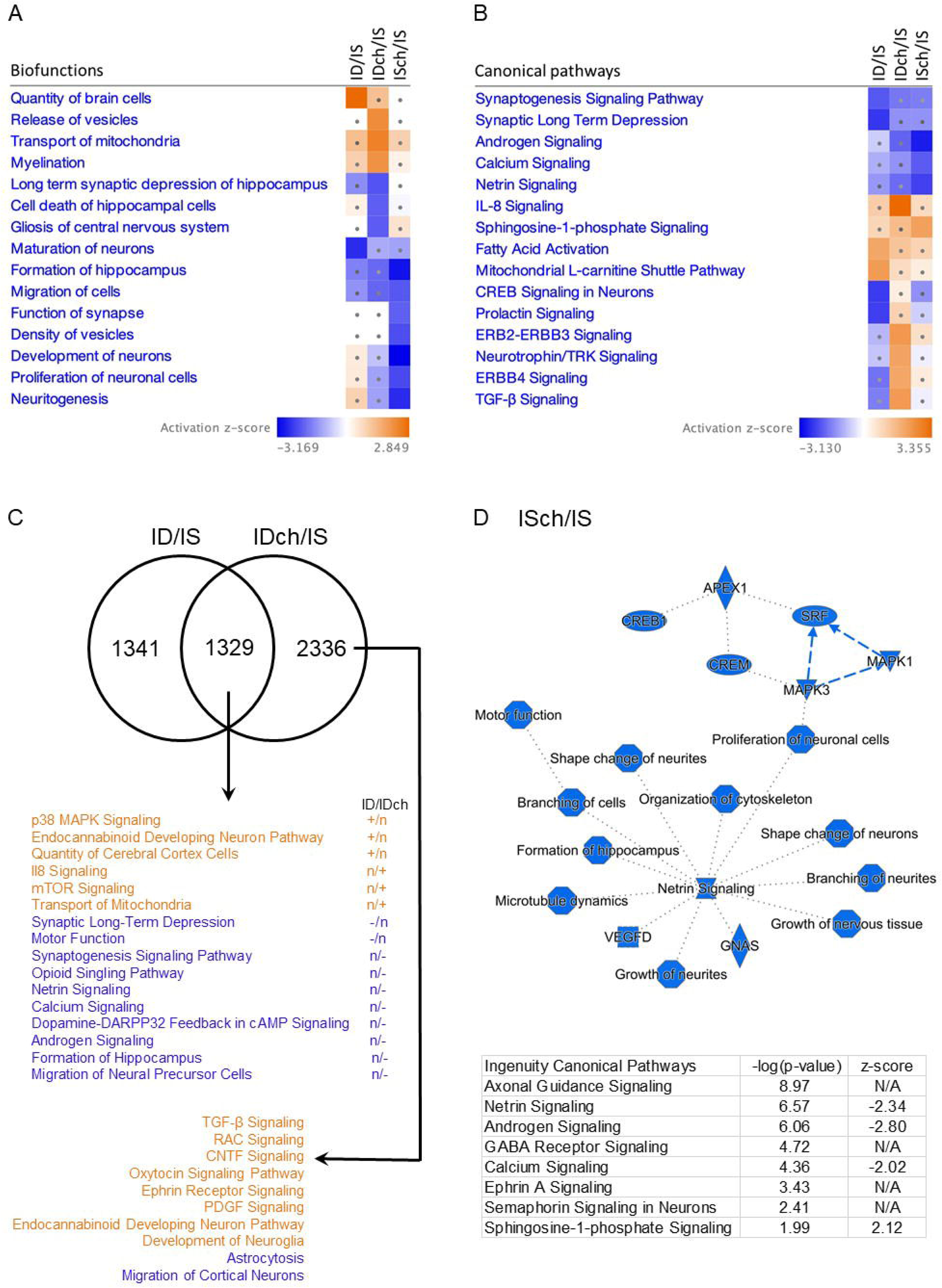
Fetal-neonatal iron deficiency and prenatal choline supplementation altered accessible chromatin (ATAC) landscape in P65 rat hippocampus. (A, B) Ingenuity-Pathway Analysis (IPA) mapped loci with differential ATAC peaks onto biofunctions (A) and canonical signaling pathways (B). Comparisons were made among iron-deficient (ID), iron-deficient choline (IDch), and iron-sufficient choline (ISch) normalized by the iron-sufficient (IS) control group. Squares with dots have absolute z-scores < 2.0, indicating non-significant findings. Blue and orange colors indicate decreased and increased chromatin accessibilities, respectively. Fewer changes were found in the ID compared to IDch and ISch groups (absolute z-scores > 2.0). Additional to reversing effects in the ID group, choline induced both decreased and increased chromatin accessibility in the IDch group. ISch group shared some similarity in ATAC pattern with ID group but also showed reduced ATAC in loci regulating synaptic structure and function. (C) Venn diagram showing overlap and non-overlapping genes modified in ID and IDch groups. +/-/n indicates increased, decreased, and normalized changes in ATAC signature. Blue/Orange indicates decreased and increased ATAC signature, respectively. (D) Comparison between ISch and IS control group showing the choline effect on ATAC landscape in adult rat hippocampus. IPA generates a graphical summary showing decreased accessibility in loci regulating hippocampal and cortical development centered on microtubule dynamics and canonical pathways showing both increased and decreased ATAC signatures. Data were selected by –log(p-value) > 1.3 and absolute z-score > 2.0.

We next used IPA to map accessible loci altered by iron deficiency or choline treatment onto known canonical pathways. Iron deficiency results in decreases accessible loci implicated in synaptic signaling, including synaptogenesis and LTD, and neurodevelopment, including CREB and prolactin signaling (z-score < -2.0, Fig. 1B, ID/IS). Conversely, iron deficiency increased chromatin accessibility of loci in energy metabolism, including fatty acid metabolism and mitochondrial function (z-score > 2.0, Fig. 1B, ID/IS). Choline treatment mitigated these effects and increased accessibility of loci critical for neuroprotection (e.g., ephrin receptor and neurotrophin/TRK signaling pathways) and neuroinflammation (IL-8 and TGF-β signaling pathways) in the IDch group (z-score > 2.0, Fig. 1B, IDch/IS). In the ISch group, choline treatment decreased accessibility of loci in the androgen, calcium and netrin signaling pathways (z-score < -2.0, Fig. 1B, ISch/IS).

To identify gene targets that are epigenetically regulated by both iron and choline, we determined the common loci, whose expression were altered by iron deficiency and choline treatment. We compared the ID/IS to IDch/IS ATAC-seq datasets and found that approximately 50% of the loci altered in the ID group were in common with those altered in the IDch group (Fig. 1C). These common loci showed changes that indicate activation of neural proliferation and development signaling pathways and inhibition of synaptic LTD and motor function in the ID group (Fig. 1C). Choline treatment not only mitigated many of the iron deficiency effects but also increased accessibility of loci regulating mitochondrial transport, IL-8 and mTOR signaling and decreased accessibility of loci regulating synaptogenesis, hippocampal formation, neuron migration, and opioid and dopamine signaling (Fig. 1C). 2,336 loci showed choline-specific effects with increased accessibility of loci in the ephrin receptor, oxytocin, and endocannabinoid signaling pathways, regulating development of neuroglia and decreased accessibility of loci regulating astrocytosis and neuronal migration (Fig. 1C).

Our previous transcriptomic study showed significant gene expression changes induced by prenatal choline supplementation in the IS adult rat hippocampus (Tran et al., 2016). Consistent with these findings, the prenatal choline treated IS (ISch) group exhibited a substantial number of changes in loci accessibility that indicate an overall reduction in gene networks regulating microtubule dynamics and neuronal development centering on reduced netrin signaling (Fig. 1D, functional network). These changes also showed a decreased accessibility of loci in the calcium and androgen, axonal guidance, GABA, and ephrin signaling pathways (Fig. 1D, bottom table).

### Prenatal choline supplementation increases enrichment of the repressive histone H3K9me3 across the adult rat hippocampal genome

To assess the long-term changes in repressive histone H3K9me3 enrichment due to either fetal-neonatal iron deficiency with or without prenatal choline supplementation, we performed a ChIP-seq analysis of the H3K9me3 profile in adult P65 rat hippocampus. Compared to the IS control group, the ID group showed a change in 1,516 differential sites that annotate to 199 loci with 89 decreased and 110 increased H3K9me3 enrichment (Table 1). Both IDch and ISch groups showed ∼23,000 differential sites that mapped onto 2,451 and 3,435 loci, respectively, based on changes at the promoters and within gene bodies (Table 1). Changes in H3K9me3 enrichment occurred predominantly in the intergenic regions (Supplemental Figure 1B).

We then functionally annotated the H3K9me3 changes using the IPA database. While iron deficiency alone resulted in no significant changes of specific biofunctions, choline treatment exhibited substantial changes in H3K9me3 enrichment. Changes in the IDch group indicate major effects with an increased enrichment in loci regulating microtubule dynamics, neurodevelopment, and neuritogenesis (z-score > 2.0, Fig. 2A, IDch/IS) as well as a decreased enrichment in loci regulating neuronal viability (z-score < -2.0, Fig. 2A, IDch/IS). The ISch group exhibited even greater H3K9me3 enrichment than the IDch group with an increased enrichment at loci regulating neural development, neuritogenesis, synaptic structures, and synaptic function (z-score > 2.0, Fig. 2A, ISch/IS) but a decreased enrichment in loci regulating behavioral deficits (z-score < -2.0, Fig. 2A, ISch/IS).

**Figure 2:**
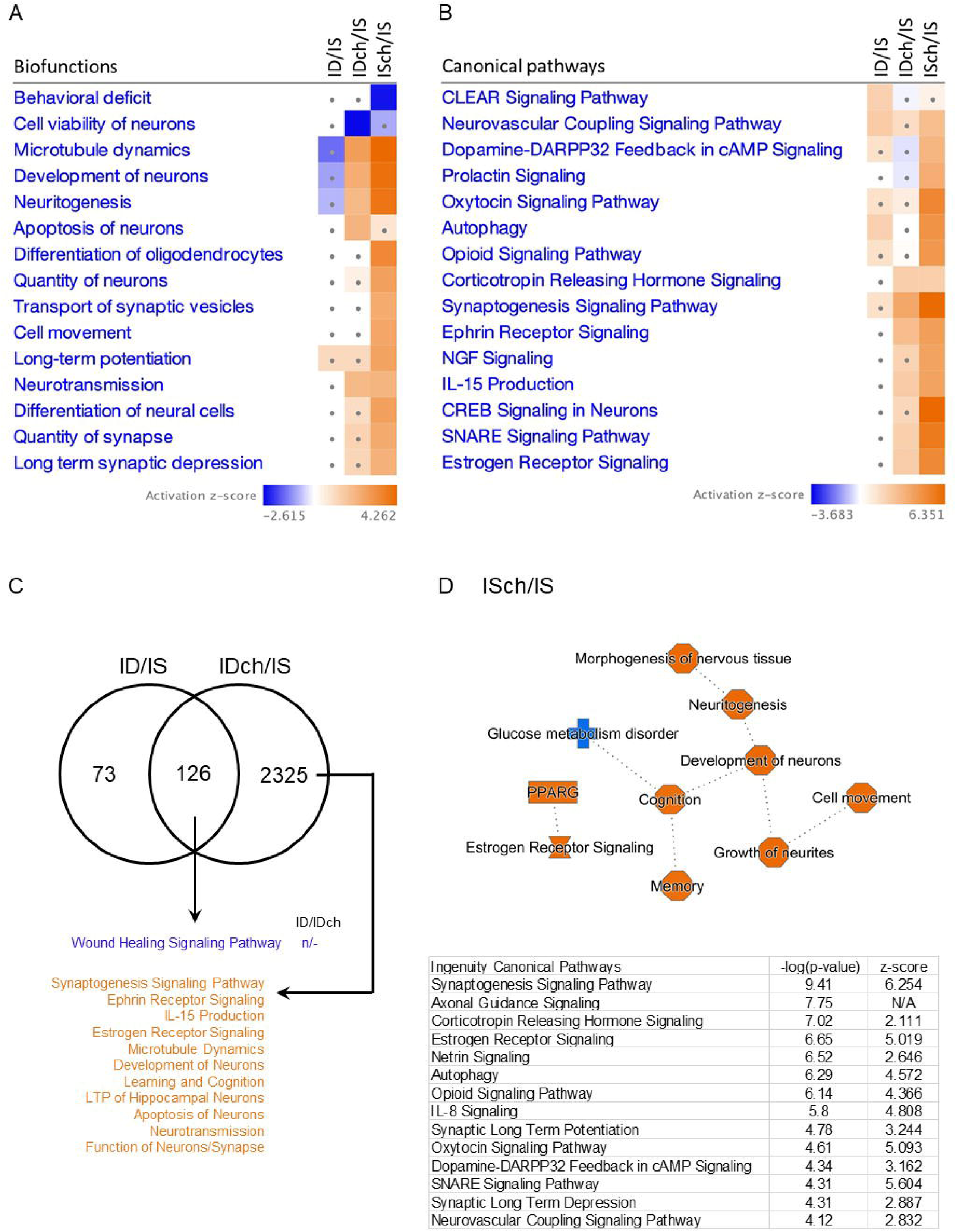
Fetal-neonatal iron deficiency and prenatal choline supplementation altered histone H3K9me3 landscape in P65 rat hippocampus. (A, B) Ingenuity-Pathway Analysis (IPA) mapped differentially-enriched H3K9me3 loci onto biofunctions (A) and canonical signaling pathways (B). Comparisons were made among iron-deficient (ID), iron-deficient choline (IDch), and iron-sufficient choline (ISch) normalized by the iron-sufficient (IS) control group. Squares with dots have absolute z-scores < 2.0. Blue and orange colors indicate decreased and increased H3K9me3 enrichment, respectively. Iron deficiency produced little changes in H3K9me3 signature (absolute z-scores < 2.0). Choline induced a significant increased H3K9me3, and even more so, enrichment in the IDch and ISch groups, respectively. (C) Venn diagram showing overlap and non-overlapping genes modified in ID and IDch groups. +/-/n indicates increased, decreased, and normalized changes in H3K9me3. Blue/Orange indicates decreased and increased H3K9me3 signature. (D) Comparison between ISch and IS control group showing the choline effect on H3K9me3 in adult rat hippocampus. IPA generates a graphical summary showing increased H3K9me3 in loci associated with cognition and estrogen receptor signaling, and canonical pathways showing significantly increased H3K9me3 enrichment. Data were selected by –log(p-value) > 1.3 and absolute z-score > 2.0.

Regarding the canonical signaling pathways, iron deficiency increased H3K9me3 enrichment at loci in synaptic function and autophagy pathways (z-score > 2.0, Fig. 2B, ID/IS). Prenatal choline treatment increased H3K9me3 enrichment at loci in the corticotropin releasing hormone (CRH), estrogen receptor (ER), and ephrin receptor signaling pathways, as well as pathways regulating neuroinflammation, synaptic structure, and synaptic function in the IDch group (z-score > 2.0, Fig. 2B, IDch/IS). The majority of these changes were choline-specific as the ISch group also showed increased H3K9me3 enrichment in many of the same pathways (Fig. 2B, ISch/IS). To further identify loci and associated biofunctions that were altered by both iron deficiency and choline, we analyzed the overlapping genes in the ID and IDch datasets. 126 loci were identified and showed a decreased enrichment in damaged tissue repair (i.e., wound healing) signaling pathway in the IDch group compared to ID group (Fig. 2C). The non-overlapping 2,325 altered genes in the IDch group showed higher H3K9me3 enrichment among loci regulating learning and cognition, microtubule dynamics, long-term potentiation (LTP) of hippocampus, and receptor (i.e., ephrin and ER) signaling pathways (z-score > 2.0, Fig. 2C,).

H3K9me3 signature changes in the ISch group indicate overall increased enrichment among loci regulating neuronal development (neuritogenesis, cell movement), and cognition (memory) as well as signaling pathways regulating synaptic transmission (synaptogenesis, synaptic LTD, LTP, SNARE, neurovascular coupling), nuclear hormone receptor signaling (ER, CRH), opioid signaling (Opioid, Dopamine-DARPP32 feedback), and neuroinflammation (IL-8, autophagy) (Fig. 2D).

### Integration of RNA-seq, ATAC-seq, and ChIP-H3K9me3-seq data revealed an increase of synaptic transmission in adult rat hippocampus due to early-life ID or prenatal choline supplementation

Congruency of transcriptomic and epigenomic signatures was compared between each treatment group (ID, IDch or ISch) and controls (IS) to further elucidate altered biofunctions and canonical pathways. There was little change with 2 loci from the ID group (Fig. 3A), 37 loci from IDch (Fig. 3B), and 107 loci from ISch group (Fig. 3C) that had changes in both chromatin marks. The 107 genes from the ISch group showed changes in expression indicating increased cell quantity and synaptic transmission (Fig. 3C). In the ID group, 22% of gene expression changes (RNA-seq) coincided with changes in chromatin accessibility (ATAC-seq). These changes indicate increased synaptic functions (synaptic transmission, depression and LTP) and decreased neural cell migration, neurite branching, and emotional behaviors (Fig. 3A). In the IDch group, 24% and 13% differentially expressed genes were associated with changes in chromatin accessibility and H3K9me3 enrichment, respectively. These changes indicate an increased in neurodevelopment, cell movement, neuroglia quantity, and cellular functions (Fig. 3B). In the ISch group, 29% and 24% of altered gene expression were related to changes in chromatin accessibility and H3K9me3 enrichment (Fig. 3C). Reductions were significant at loci associated with neural and axonal development, while increases were significant among loci associated with cell migration, synaptic transmission, and memory (Fig. 3C).

**Figure 3:**
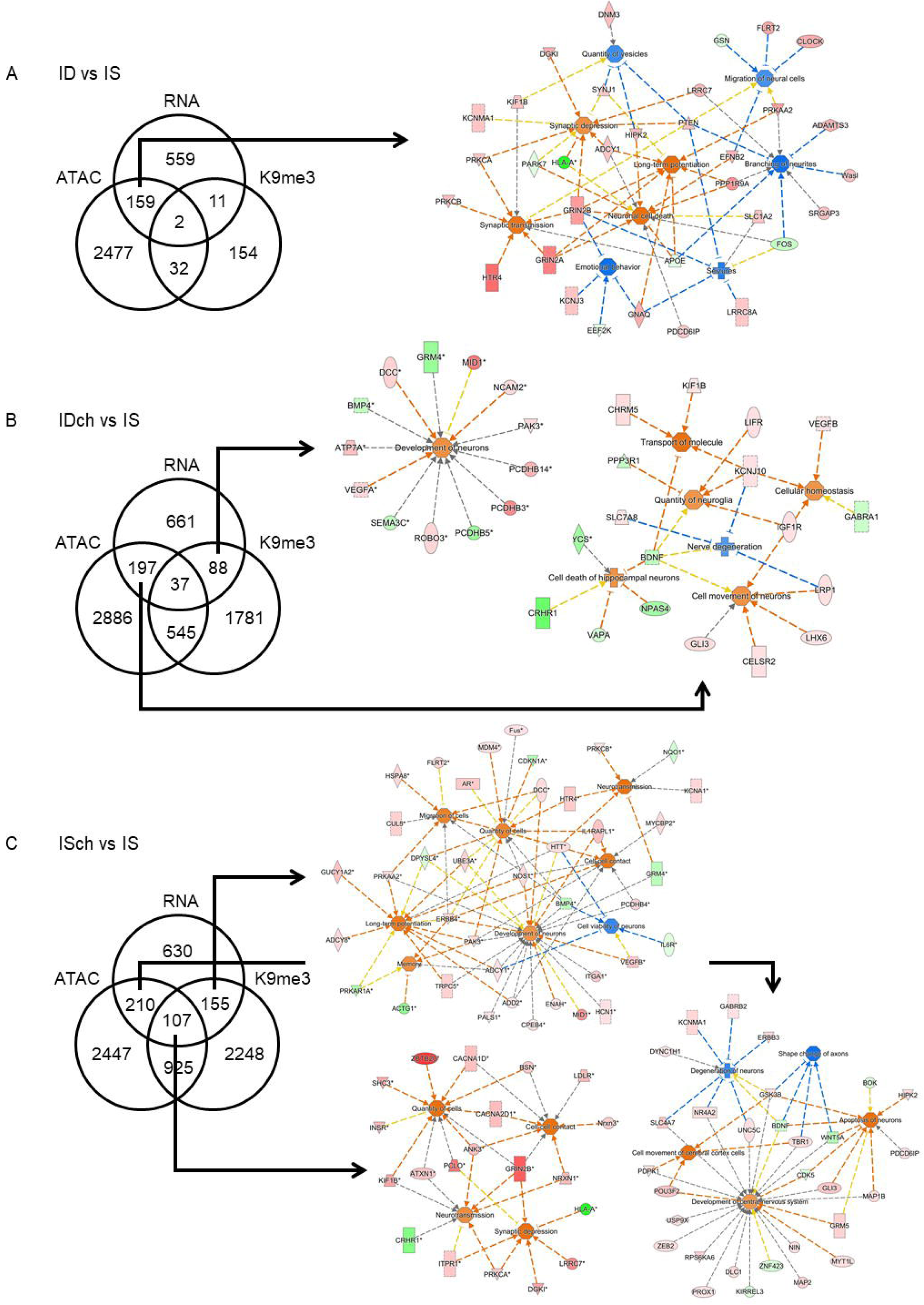
Integration of P65 hippocampal RNA-seq, ATAC-seq, and ChIP-H3K9me3-seq data. (A) ID and IS comparison showing 161 of 731 (22.0%) overlapping loci between ATAC and RNA datasets. These differentially expressed genes indicate increased synaptic transmission, synaptic depression and potentiation, and cell death accompanied by decreased branching of neurites. (B) IDch vs IS comparison showing 125 loci changing in H3K9me3 signature, which suggests a potential activation in neuron development. 234 loci showed changes in chromatin accessibility that indicate an increase in cellular function. (C) ISch and IS comparison showing 107 overlapping loci among all datasets, which indicate higher synaptic signaling and cell proliferation. 262 loci showed changes in H3K9me3 signature, suggesting impaired neuronal viability and increased cell signaling. 317 loci showed changes in accessibility that indicate reduced neurodegeneration accompanied by increased CNS development.

To provide a granular analysis of chromatin changes induced by ID or choline treatment with respect to accessibility and H3K9me3-silencing signatures, we examined the aligned sequences using IGV tools (Broad Institute). The data revealed a specific pattern of changes with both significant (e.g., chromosome 10) and little (e.g., chromosome 12) modification among different chromosomes (Fig. 4A, Supplemental Figures 2 and 3). ATAC and H3K9me3 peaks showed overlap mainly in the intergenic regions but not at the promoters or gene bodies among all four experimental groups (Table 2). Further analysis of the nearest genes to these overlapping regions showed that the 91 common genes among IS, ID and IDch groups implicate a gene network centering on BDNF, PSEN1, and APP (Fig. 4B) and regulate nervous system development. Choline-specific effect highlighted 142 genes associated with a gene network that centers on SNCA, HTT, and TCF7L2, which are associated with neuropathology such as Parkinson’s and Huntington’s diseases (Fig. 4C).

**Figure 4:**
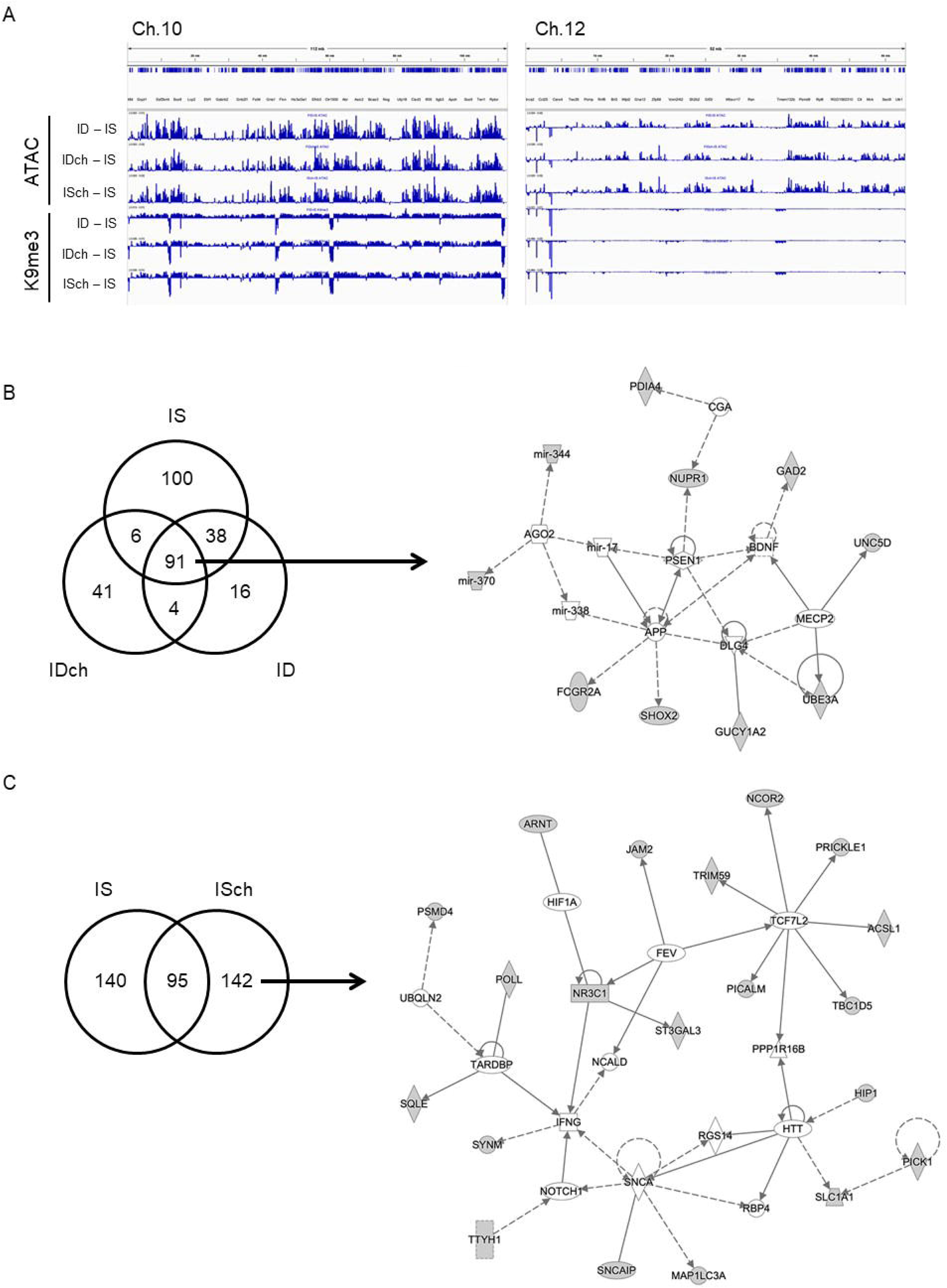
Alignments of ATAC-seq and H3K9me3-seq peaks. (A) Aligned and annotated Tn5 transposase (ATAC) and H3K9me3 coverage plots along a 112-mb and 52-mb region of Chromosome 10 and 12. (B) Overlapping ATAC and H3K9me3 peaks among IS, ID, and IDch groups localize to the intergenic regions of 91 nearby genes, which mapped onto a gene network centering on the Brain-derived neurotrophic factor (BDNF), Presenilin-1 (PSEN1), and Amyloid precursor protein (APP) that regulate nervous system development. (C) Overlapping ATAC and H3K9me3 peaks between IS and ISch groups showed 142 choline-specific effect genes that mapped onto a gene network centering on alpha-synuclein (SNCA), Huntingtin (HTT), and transcritipion factor 7 like 2 (TCF7L2), which are associated with neurological diseases.

**Table 2:** Overlapping regions between ATAC and H3K9me3 peaks. The overlapping peaks between the two datasets were primarily located in the intergenic regions.

| *Treatment Group* | *# Overlapping Regions* | *# Nearest Loci* | *IPA-Annotated Diseases and Functions* |
| --- | --- | --- | --- |
| *IS* | 868 | 235 | Development of CNS, Long-Term Potentiation (LTP) |
| *ISch* | 593 | 237 | Synaptic transmission of cerebrocortical cells, LTP of hippocampal neurons |
| *ID* | 551 | 149 | Differentiation of neurons, Migration of cells, LTP |
| *IDch* | 563 | 142 | Cerebral disorder, Tauopathy, LTP, Migration of neuroglia |

To further illustrate the chromatin changes, differential peaks at bone morphogenetic protein 3 (*Bmp3*), tankyrase (*Tnks*), and iron responsive element binding protein 2 (*Ireb2*) were compared among treatment groups (Fig. 5A). Compared to the IS control group, these genes exhibited increased ATAC peaks in the ID group (all 3 genes) and ISch group (*Bmp3* and *Tnks*), and little change in the IDch group (Fig. 5A, boxed areas). We further identified DNA sequences at these specific sites with potential transcription factor binding, including MYC, upstream stimulating factor (USF), CREB, CCAAT/enhancer binding protein beta (CEBPB), signal transducer and activator of transcription (STAT), NF-E2 related factor 2 (NRF2), and ETS transcription factor ELK1 (ELK1) (Fig. 5A, bottom text). ChIP-H3K9me3 results at these loci were not above background noise. These chromatin changes were positively correlated with changes in gene expression, where both ID and ISch groups showed a higher expression level compared to IS control group (Fig. 5B).

**Figure 5:**
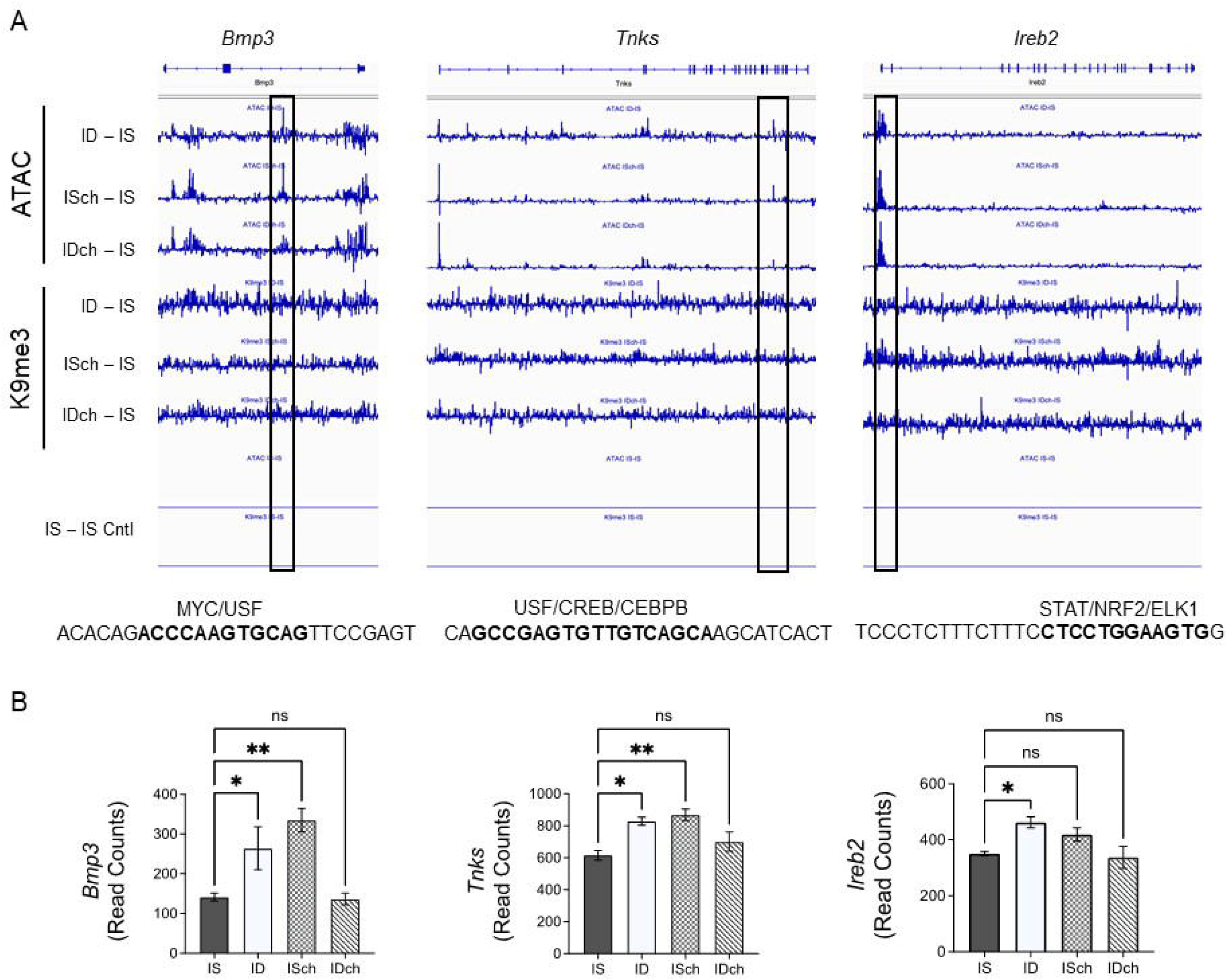
Alignments of ATAC and H3K9me3 peaks. (A) Peaks aligned to Bmp3, Tnks, and Ireb2 loci showing differential effects by iron deficiency and choline. Brackets depict regions with differential peaks among ID, ISch, and IDch groups and potential enhancers with specific transcription factor binding sites. (B) Graphs showing transcript counts of Bmp3, Tnks, and Ireb2 generated from RNA-seq data. Values are mean +/- SEM; asterisks denote *p<0.05 and **p<0.01.

## DISCUSION

Early-life iron deficiency produces persistent and widespread hippocampal gene dysregulation in preclinical models (Tran et al., 2009, 2016). We previously demonstrated changes of DNA methylation in the developing iron-deficient hippocampus (Lien et al., 2019) and chromatin methylation at the *Bdnf* gene in the formerly iron-deficient hippocampus (Tran et al., 2016), providing evidence for epigenetic modifications due to iron deficiency. To further establish that early-life iron deficiency induces long-term epigenetic changes in the adult hippocampus, the present study assessed and integrated signatures of accessible chromatin modifications (ATAC/active vs. H3K9me3/repressive) and transcriptomic expression from the formerly ID rat hippocampus. The findings reveal that iron deficiency reduced chromatin accessibility accounting for about 22% of gene expression changes, particularly among loci that regulate neurite growth and synaptic transmission, and had little (∼1.7%) effect on the repressive histone H3K9me3 mark, indicating an incomplete account for the gene expression changes and involvement of other epigenetic regulators. Thus, integration of additional epigenetic marks (e.g., H3K27me3, H3K4me3, 5mC, and 5hmC) is needed in future studies to more fully elucidate mechanisms contributing to the long-term gene dysregulation in the developing central nervous system (Maze et al., 2014; Feng et al., 2015). It is also possible that the small overlap of genes among the three analyses were resulted from variations in biological samples as different rats were used for each type of analyses. This possibility could be addressed in future studies that employ multi-omics analyses of single nucleus sequencing approach (Habib et al., 2016). As such, these findings are consistent with findings in the human studies that demonstrate limited correlation when assessing a single epigenetic mark (Ouyang et al., 2009). In addition, the effects of prenatal choline supplementation were also evaluated given its initial identification as a potential adjunctive treatment for fetal-neonatal iron deficiency (Kennedy et al., 2016, 2018). Choline partially rescued the effects of iron deficiency, accounting for 24% (vs 22%) of gene expression changes. Prenatal choline supplementation produced an even greater effect on the H3K9me3 signature in the IS group, specifically among loci regulating neurotransmission and opioid signaling. Collectively, these findings reveal a lasting consequence of early-life iron deficiency and choline supplementation on the adult rat hippocampal epigenome.

Iron deficiency resulted in more loci showing increased than decreased accessibility, consistent with our transcriptomic data. These findings support the conclusion that changes in ATAC account for a part of the long-term epigenetic modifications induced by early-life iron deficiency (Liao et al., 2020). Nevertheless, it is not surprising that the reduced accessibility occurred among loci regulating maturation of neurons and development of hippocampal-cerebrocortical circuit, which may explain the impaired neurocognitive function exhibited by formerly ID animals (Kennedy et al., 2014, 2018). Our group recently reported the effect of iron deficiency on neuroinflammation (Singh et al., 2021). Given the important role of microglia in neural development and function (Szepesi et al., 2018), the long-term decrease of neuroinflammation signaling pathways could contribute to the abnormal cognition and emotional behavior observed in the formerly ID adult rats (Carlson et al., 2007; Pisansky et al., 2013).

The established beneficial effects of prenatal choline supplementation demonstrated by others (Zeisel, 2004; Mellott et al., 2017; Chin et al., 2019; Agam et al., 2020) and by us in this model of fetal-neonatal iron deficiency (Kennedy et al., 2016; Tran et al., 2016) and choline being a substrate for epigenetic modifications (Zeisel, 2017) justified the rationale for assessing the long-term neuroepigenomic effects. A major finding was the long-term effects of choline on both ATAC and H3K9me3 signatures in the hippocampus of the formerly ID and IS control groups. In the ID group, choline supplementation normalized iron deficiency-reduced accessibility of genes that regulate neuronal maturation, synaptogenesis, LTD, and CREB signaling pathways as well as promoted accessibility of genes regulating myelination, mitochondrial transport, synaptic function, and ephrin and neurotrophin signaling pathways. These effects could account for the partial recovery of hippocampal-dependent learning and memory function in IDch adult rats (Kennedy et al., 2016). While these long-term epigenetic changes were found in the hippocampus, it is possible that they also occurred in the cerebral cortex and cerebellum, consistent with a long, developmentally sensitive window in these structures as they have a protracted developmental timeline (Andersen, 2003). Future studies could investigate the epigenomic effects of iron deficiency on other developing brain regions as we have begun in a previous study (Barks et al., 2022).

A novel and unique finding in prenatal choline supplementation to the IS group (ISch) was the increased H3K9me3 mark among loci regulating opioid and dopamine-DARPP32 signaling pathways, suggesting that choline can influence the reward circuitry potentially involving the hippocampal-cortical-nucleus accumbens connectivity (Le Merre et al., 2018; LeGates et al., 2018). The finding raises possible concerns as to whether prenatal choline supplementation may affect drug-seeking behaviors in adulthood given the chromatin changes in loci implicating altered balance among dopaminergic, glutamatergic, and GABAergic synaptic transmission (Tang and Dani, 2009; Enoch et al., 2012; Castilla-Ortega et al., 2016; Márquez et al., 2016). Thus, the use of choline as a therapeutic agent needs more studies to determine the appropriate dose and duration during gestation to maximize the beneficial effects while minimize potential negative impacts.

The effects of ID or choline on ATAC and H3K9me3 enrichment were largely distinct with limited overlapping regions, indicating that the majority of peaks between ATAC and H3K9me3 occurred at non-overlapping regions. However, the overlapping regions of ATAC and H3K9me3 peaks altered by early-life ID locate near a gene network that regulates well-documented ID-altered factors, including BDNF, APP and PSEN1 (Carlson et al., 2007; Tran et al., 2009). The findings highlight convergent epigenetic modifications and reveal potential long-distance regulatory elements for specific gene networks known to be altered by early-life iron deficiency. Likewise, prenatal choline produced convergent epigenetic modifications near a specific gene network regulating factors that contribute to neurological diseases, including SNCA, HTT, and NR3C1. Thus, it is important to determine the long-term neurological effects of prenatal choline supplementation to fully evaluate its beneficial effects beyond cognitive function.

In conclusion, early-life iron deficiency produces lasting and widespread changes in the rat hippocampal epigenome, providing molecular bases for the long-term gene dysregulations. This study provides evidence that changes in chromatin accessibility could partially account for gene dysregulation. The mechanisms underlying changes in chromatin landscape were unlikely involved changes in H3K9me3 mark, thereby excluding the possible involvement of iron-dependent histone demethylases (e.g., KDM4) and methyltransferases (e.g., G9A) that target histone H3K9. Conversely, choline-mediated epigenetic changes in the rat hippocampus involved modification of the H3K9me3 landscape, implicating H3K9 methyltransferases and demethylases. Since epigenetic regulation involves complex interactions among iron-dependent and -independent factors, this study provides a foundation upon which additional studies can build to fully elucidate mechanisms and identify targets for therapeutic development in the prevention of long-term adverse outcomes of early-life exposures.

## Conflict of interest statement

All authors have no conflict of interest to disclose.

## Author’s contribution

PVT and MKG developed conceptual framework; SXL and PVT conducted experiments and analyzed results; AR and LS performed bioinformatic analyses; SXL and PVT composed the manuscript; All authors reviewed, edited, and approved the manuscript.

## Acknowledgments

We thank Dr. Juan E. Abrahante Lloréns for help with RNA-seq data.

## Funding sources

This work was funded by NIH R01NS099178 to PVT and R01HD29421 to MKG

## Notes

### Competing Interest Statement

The authors have declared no competing interest.

## References

Agam G, Taylor Z, Vainer E, Golan HM (2020) The influence of choline treatment on behavioral and neurochemical autistic-like phenotype in Mthfr-deficient mice. Transl Psychiatry 2020 101 10:1–14 Available at: https://www.nature.com/articles/s41398-020-01002-1 [Accessed August 17, 2021].

Andersen SL (2003) Trajectories of brain development: point of vulnerability or window of opportunity? Neurosci Biobehav Rev 27:3–18.

Barks A, Beeson MM, Hallstrom TC, Georgieff MK, Tran P V. (2022) Developmental Iron Deficiency Dysregulates TET Activity and DNA Hydroxymethylation in the Rat Hippocampus and Cerebellum. Dev Neurosci 44:80–90 Available at: https://www.karger.com/Article/FullText/521704 [Accessed June 6, 2022].

Barks A, Fretham SJB, Georgieff MK, Tran P V. (2018) Early-Life Neuronal-Specific Iron Deficiency Alters the Adult Mouse Hippocampal Transcriptome. J Nutr 148:1521–1528.

Barks A, Hall AM, Tran P V., Georgieff MK (2019) Iron as a model nutrient for understanding the nutritional origins of neuropsychiatric disease. Pediatr Res 85:176–182 Available at: https://pubmed.ncbi.nlm.nih.gov/30341413/ [Accessed March 21, 2021].

Buenrostro J, Wu B, Chang H, Greenleaf W (2015) ATAC-seq: A Method for Assaying Chromatin Accessibility Genome-Wide. Curr Protoc Mol Biol 109:21.29.1 Available at: /pmc/articles/PMC4374986/ [Accessed September 20, 2021].

Carlson ES, Stead JDH, Neal CR, Petryk A, Georgieff MK (2007) Perinatal iron deficiency results in altered developmental expression of genes mediating energy metabolism and neuronal morphogenesis in hippocampus. Hippocampus 17:679–691 Available at: https://pubmed.ncbi.nlm.nih.gov/17546681/ [Accessed March 21, 2021].

Castilla-Ortega E, Serrano A, Blanco E, Araos P, Suárez J, Pavón FJ, Rodríguez de Fonseca F, Santín LJ (2016) A place for the hippocampus in the cocaine addiction circuit: Potential roles for adult hippocampal neurogenesis. Neurosci Biobehav Rev 66:15–32.

Chin EWM, Lim WM, Ma D, Rosales FJ, Goh ELK (2019) Choline Rescues Behavioural Deficits in a Mouse Model of Rett Syndrome by Modulating Neuronal Plasticity. Mol Neurobiol 56:3882–3896 Available at: https://pubmed.ncbi.nlm.nih.gov/30220058/ [Accessed August 17, 2021].

Cooney CA, Dave AA, Wolff GL (2002) Maternal methyl supplements in mice affect epigenetic variation and DNA methylation of offspring. J Nutr 132:2393S–2400S.

Corces MR et al. (2017) An improved ATAC-seq protocol reduces background and enables interrogation of frozen tissues. Nat Methods 2017 1410 14:959–962 Available at: https://www.nature.com/articles/nmeth.4396 [Accessed August 17, 2021].

Davison JM, Mellott TJ, Kovacheva VP, Blusztajn JK (2009) Gestational choline supply regulates methylation of histone H3, expression of histone methyltransferases G9a (Kmt1c) and Suv39h1 (Kmt1a), and DNA methylation of their genes in rat fetal liver and brain. J Biol Chem 284:1982–1989 Available at: /pmc/articles/PMC2629111/ [Accessed March 21, 2021].

Enoch M-A, Zhou Z, Kimura M, Mash DC, Yuan Q, Goldman D (2012) GABAergic Gene Expression in Postmortem Hippocampus from Alcoholics and Cocaine Addicts; Corresponding Findings in Alcohol-Naïve P and NP Rats. PLoS One 7:e29369 Available at: https://journals.plos.org/plosone/article?id=10.1371/journal.pone.0029369 [Accessed July 29, 2021].

Feng J et al. (2015) Role of Tet1 and 5-hydroxymethylcytosine in cocaine action. Nat Neurosci 2015 184 18:536–544 Available at: https://www.nature.com/articles/nn.3976 [Accessed September 20, 2021].

Fisher MC, Zeisel SH, Mar MH, Sadler TW (2002) Perturbations in choline metabolism cause neural tube defects in mouse embryos in vitro. FASEB J 16:619–621 Available at: https://pubmed.ncbi.nlm.nih.gov/11919173/ [Accessed March 21, 2021].

Fretham SJB, Carlson ES, Georgieff MK (2011) The role of iron in learning and memory. Adv Nutr 2:112–121 Available at: /pmc/articles/PMC3065765/ [Accessed March 21, 2021].

Fukumitsu K, Hatsukano T, Yoshimura A, Heuser J, Fujishima K, Kengaku M (2016) Mitochondrial fission protein Drp1 regulates mitochondrial transport and dendritic arborization in cerebellar Purkinje cells. Mol Cell Neurosci 71:56–65 Available at: https://pubmed.ncbi.nlm.nih.gov/26689905/ [Accessed March 21, 2021].

Glenn MJ, Kirby ED, Gibson EM, Wong-Goodrich SJ, Mellott TJ, Blusztajn JK, Williams CL (2008) Age-related declines in exploratory behavior and markers of hippocampal plasticity are attenuated by prenatal choline supplementation in rats. Brain Res 1237:110–123 Available at: /pmc/articles/PMC2677022/ [Accessed March 21, 2021].

Habib N, Li Y, Heidenreich M, Swiech L, Avraham-Davidi I, Trombetta JJ, Hession C, Zhang F, Regev A (2016) Div-Seq: Single-nucleus RNA-Seq reveals dynamics of rare adult newborn neurons. Science 353:925–928 Available at: https://pubmed.ncbi.nlm.nih.gov/27471252/ [Accessed June 6, 2022].

Jirtle RL (2014) The Agouti mouse: A biosensor for environmental epigenomics studies investigating the developmental origins of health and disease. Epigenomics 6:447–450 Available at: https://www.futuremedicine.com/doi/full/10.2217/epi.14.58 [Accessed May 16, 2022].

Kennedy BC, Dimova JG, Siddappa AJM, Tran P V, Gewirtz JC, Georgieff MK (2014) Prenatal choline supplementation ameliorates the long-term neurobehavioral effects of fetal-neonatal iron deficiency in rats. J Nutr 144:1858–1865 Available at: http://www.ncbi.nlm.nih.gov/pubmed/25332485 [Accessed October 7, 2019].

Kennedy BC, Tran P V., Kohli M, Maertens JJ, Gewirtz JC, Georgieff MK (2018) Beneficial effects of postnatal choline supplementation on long-Term neurocognitive deficit resulting from fetal-Neonatal iron deficiency. Behav Brain Res 336:40–43 Available at: https://pubmed.ncbi.nlm.nih.gov/28811181/ [Accessed March 21, 2021].

Kennedy BC, Wallin DJ, Tran P V., Georgieff MK (2016) Long-term brain and behavioral consequences of early-life iron deficiency. In: Fetal Development: Research on Brain and Behavior, Environmental Influences, and Emerging Technologies, pp 295–316 Available at: https://academic.oup.com/nutritionreviews/article-lookup/doi/10.1111/j.1753-4887.2011.00432.x [Accessed October 7, 2019].

Kim D, Paggi JM, Park C, Bennett C, Salzberg SL (2019) Graph-Based Genome Alignment and Genotyping with HISAT2 andHISAT-genotype. Nat Biotechnol 37:907 Available at: /pmc/articles/PMC7605509/ [Accessed August 17, 2021].

Krzysztof Blusztajn J, J. Mellott T (2012) Choline Nutrition Programs Brain Development Via DNA and Histone Methylation. Cent Nerv Syst Agents Med Chem 12:82–94 Available at: https://pubmed.ncbi.nlm.nih.gov/22483275/ [Accessed March 21, 2021].

Kuzawa CW (1998) Adipose Tissue in Human Infancy and Childhood: An Evolutionary Perspective. Yearb Phys Anthropol 41:177–209 Available at: https://pubmed.ncbi.nlm.nih.gov/9881526/ [Accessed March 21, 2021].

Le Merre P, Esmaeili V, Charrière E, Galan K, Salin PA, Petersen CCH, Crochet S (2018) Reward-Based Learning Drives Rapid Sensory Signals in Medial Prefrontal Cortex and Dorsal Hippocampus Necessary for Goal-Directed Behavior. Neuron 97:83–91.e5.

LeGates TA, Kvarta MD, Tooley JR, Francis TC, Lobo MK, Creed MC, Thompson SM (2018) Reward behaviour is regulated by the strength of hippocampus–nucleus accumbens synapses. Nat 2018 5647735 564:258–262 Available at: https://www.nature.com/articles/s41586-018-0740-8 [Accessed June 6, 2022].

Li Z, Okamoto KI, Hayashi Y, Sheng M (2004) The importance of dendritic mitochondria in the morphogenesis and plasticity of spines and synapses. Cell 119:873–887 Available at: https://pubmed.ncbi.nlm.nih.gov/15607982/ [Accessed March 21, 2021].

Liao R, Zheng Y, Liu X, Zhang Y, Seim G, Tanimura N, Wilson GM, Hematti P, Coon JJ, Fan J, Xu J, Keles S, Bresnick EH (2020) Discovering How Heme Controls Genome Function Through Heme-omics. Cell Rep 31:107832.

Lien YC, Condon DE, Georgieff MK, Simmons RA, Tran P V. (2019) Dysregulation of neuronal genes by fetal-neonatal iron deficiency anemia is associated with altered DNA methylation in the rat hippocampus. Nutrients 11:1191 Available at: https://pubmed.ncbi.nlm.nih.gov/31137889/ [Accessed March 21, 2021].

Liu SX, Barks AK, Lunos S, Gewirtz JC, Georgieff MK, Tran P V. (2021) Prenatal Iron Deficiency and Choline Supplementation Interact to Epigenetically Regulate Jarid1b and Bdnf in the Rat Hippocampus into Adulthood. Nutrients 13 Available at: https://pubmed.ncbi.nlm.nih.gov/34960080/ [Accessed March 30, 2022].

Lozoff B, Jimenez E, Hagen J, Mollen E, Wolf AW (2000) Poorer behavioral and developmental outcome more than 10 years after treatment for iron deficiency in infancy. Pediatrics 105:e51 Available at: https://pubmed.ncbi.nlm.nih.gov/10742372/ [Accessed March 21, 2021].

Lozoff B, Smith JB, Kaciroti N, Clark KM, Guevara S, Jimenez E (2013) Functional significance of early-life iron deficiency: Outcomes at 25 years. J Pediatr 163:1260–1266.

Márquez J, Campos-Sandoval JA, Peñalver A, Matés JM, Segura JA, Blanco E, Alonso FJ, de Fonseca FR (2016) Glutamate and Brain Glutaminases in Drug Addiction. Neurochem Res 2016 423 42:846–857 Available at: https://link.springer.com/article/10.1007/s11064-016-2137-0 [Accessed August 17, 2021].

Maze I, Shen L, Zhang B, Garcia BA, Shao N, Mitchell A, Sun H, Akbarian S, Allis CD, Nestler EJ (2014) Analytical tools and current challenges in the modern era of neuroepigenomics. Nat Neurosci 2014 1711 17:1476–1490 Available at: https://www.nature.com/articles/nn.3816 [Accessed September 20, 2021].

Mellott TJ, Huleatt OM, Shade BN, Pender SM, Liu YB, Slack BE, Blusztajn JK (2017) Perinatal Choline Supplementation Reduces Amyloidosis and Increases Choline Acetyltransferase Expression in the Hippocampus of the APPswePS1dE9 Alzheimer’s Disease Model Mice. PLoS One 12 Available at: https://pubmed.ncbi.nlm.nih.gov/28103298/ [Accessed August 17, 2021].

Ouyang Z, Zhou Q, Wong WH (2009) ChIP-Seq of transcription factors predicts absolute and differential gene expression in embryonic stem cells. Proc Natl Acad Sci U S A 106:21521 Available at: /pmc/articles/PMC2789751/ [Accessed June 6, 2022].

Pisansky MT, Wickham RJ, Su J, Fretham S, Yuan LL, Sun M, Gewirtz JC, Georgieff MK (2013) Iron deficiency with or without anemia impairs prepulse inhibition of the startle reflex. Hippocampus 23:952–962 Available at: /pmc/articles/PMC3888485/ [Accessed March 21, 2021].

Schachtschneider KM, Liu Y, Rund LA, Madsen O, Johnson RW, Groenen MAM, Schook LB (2016) Impact of neonatal iron deficiency on hippocampal DNA methylation and gene transcription in a porcine biomedical model of cognitive development. BMC Genomics 17:1–14 Available at: /pmc/articles/PMC5094146/ [Accessed August 30, 2021].

Shen L, Shao N-Y, Liu X, Maze I, Feng J, Nestler EJ (2013) diffReps: Detecting Differential Chromatin Modification Sites from ChIP-seq Data with Biological Replicates. PLoS One 8:e65598 Available at: https://journals.plos.org/plosone/article?id=10.1371/journal.pone.0065598 [Accessed August 17, 2021].

Singh G, Segura BJ, Georgieff MK, Gisslen T (2021) Fetal inflammation induces acute immune tolerance in the neonatal rat hippocampus. J Neuroinflammation 18 Available at: /pmc/articles/PMC7953777/ [Accessed August 17, 2021].

Szepesi Z, Manouchehrian O, Bachiller S, Deierborg T (2018) Bidirectional Microglia–Neuron Communication in Health and Disease. Front Cell Neurosci 0:323.

Tang J, Dani JA (2009) Dopamine Enables In Vivo Synaptic Plasticity Associated with the Addictive Drug Nicotine. Neuron 63:673–682.

Tran P V., Fretham SJB, Carlson ES, Georgieff MK (2009) Long-term reduction of hippocampal brain-derived neurotrophic factor activity after fetal-neonatal iron deficiency in adult rats. Pediatr Res 65:493–498 Available at: https://pubmed.ncbi.nlm.nih.gov/19190544/ [Accessed March 21, 2021].

Tran P V., Fretham SJB, Wobken J, Miller BS, Georgieff MK (2012) Gestational-neonatal iron deficiency suppresses and iron treatment reactivates IGF signaling in developing rat hippocampus. Am J Physiol - Endocrinol Metab 302:316–324 Available at: /pmc/articles/PMC3287363/ [Accessed March 21, 2021].

Tran P V., Kennedy BC, Lien Y-C, Simmons RA, Georgieff MK (2015) Fetal iron deficiency induces chromatin remodeling at the Bdnf locus in adult rat hippocampus. Am J Physiol Integr Comp Physiol 308:R276–R282 Available at: https://www.physiology.org/doi/10.1152/ajpregu.00429.2014 [Accessed March 16, 2021].

Tran P V., Kennedy BC, Pisansky MT, Won KJ, Gewirtz JC, Simmons RA, Georgieff MK (2016) Prenatal choline supplementation diminishes early-life iron deficiency-induced reprogramming of molecular networks associated with behavioral abnormalities in the adult rat hippocampus. J Nutr 146:484–493 Available at: https://pubmed.ncbi.nlm.nih.gov/26865644/ [Accessed March 21, 2021].

Tsunoda T, Takagi T (1999) Estimating transcription factor bindability on DNA. Bioinformatics 15:622–630 Available at: https://pubmed.ncbi.nlm.nih.gov/10487870/ [Accessed August 17, 2021].

Wong-Goodrich SJE, Glenn MJ, Mellott TJ, Blusztajn JK, Meck WH, Williams CL (2008) Spatial memory and hippocampal plasticity are differentially sensitive to the availability of choline in adulthood as a function of choline supply in utero. Brain Res 1237:153–166 Available at: /pmc/articles/PMC2674276/ [Accessed March 21, 2021].

Wozniak JR, Fuglestad AJ, Eckerle JK, Kroupina MG, Miller NC, Boys CJ, Brearley AM, Fink BA, Hoecker HL, Zeisel SH, Georgieff MK (2013) Choline supplementation in children with Fetal Alcohol Spectrum Disorders (FASD) has high feasibility and tolerability. Nutr Res 33:897–904 Available at: /pmc/articles/PMC3815698/ [Accessed July 16, 2021].

Zeisel S (2017) Choline, Other Methyl-Donors and Epigenetics. Nutr 2017, Vol 9, Page 445 9:445 Available at: https://www.mdpi.com/2072-6643/9/5/445/htm [Accessed August 30, 2021].

Zeisel SH (2004) Nutritional Importance of Choline for Brain Development. J Am Coll Nutr 23:621S–626S Available at: http://www.tandfonline.com/doi/abs/10.1080/07315724.2004.10719433 [Accessed March 25, 2021].

Zhang Y, Liu T, Meyer CA, Eeckhoute J, Johnson DS, Bernstein BE, Nusbaum C, Myers RM, Brown M, Li W, Liu XS (2008) Model-based Analysis of ChIP-Seq (MACS). Genome Biol 2008 99 9:1–9 Available at: https://genomebiology.biomedcentral.com/articles/10.1186/gb-2008-9-9-r137 [Accessed August 17, 2021].

