## Supplemental Figures 1-3 for "ATAC and histone H3K9me3 landscapes revealed the altered epigenome by fetal-neonatal iron deficiency in the adult male rat hippocampus"

### Slide 1
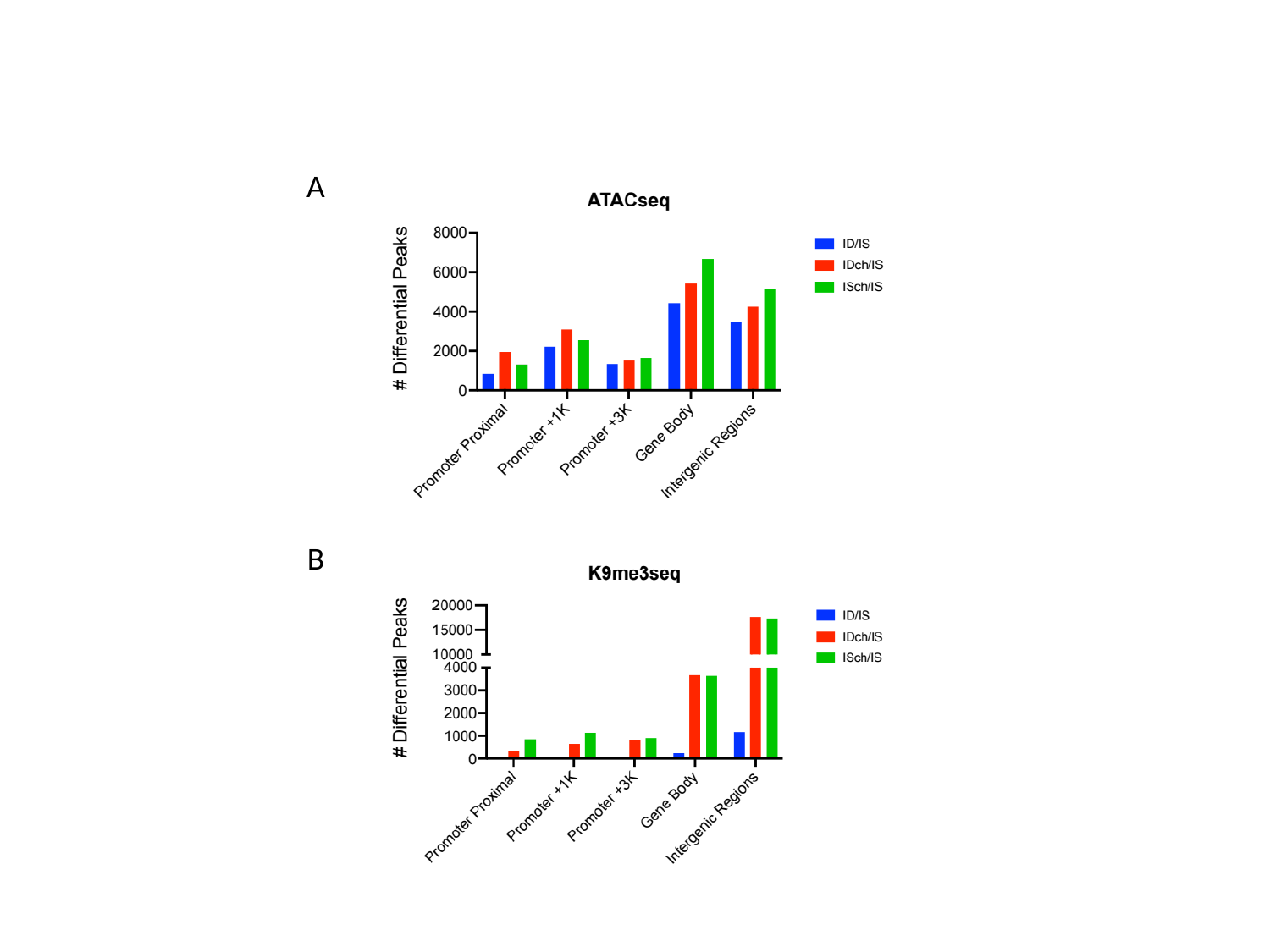

A
B

### Slide 2
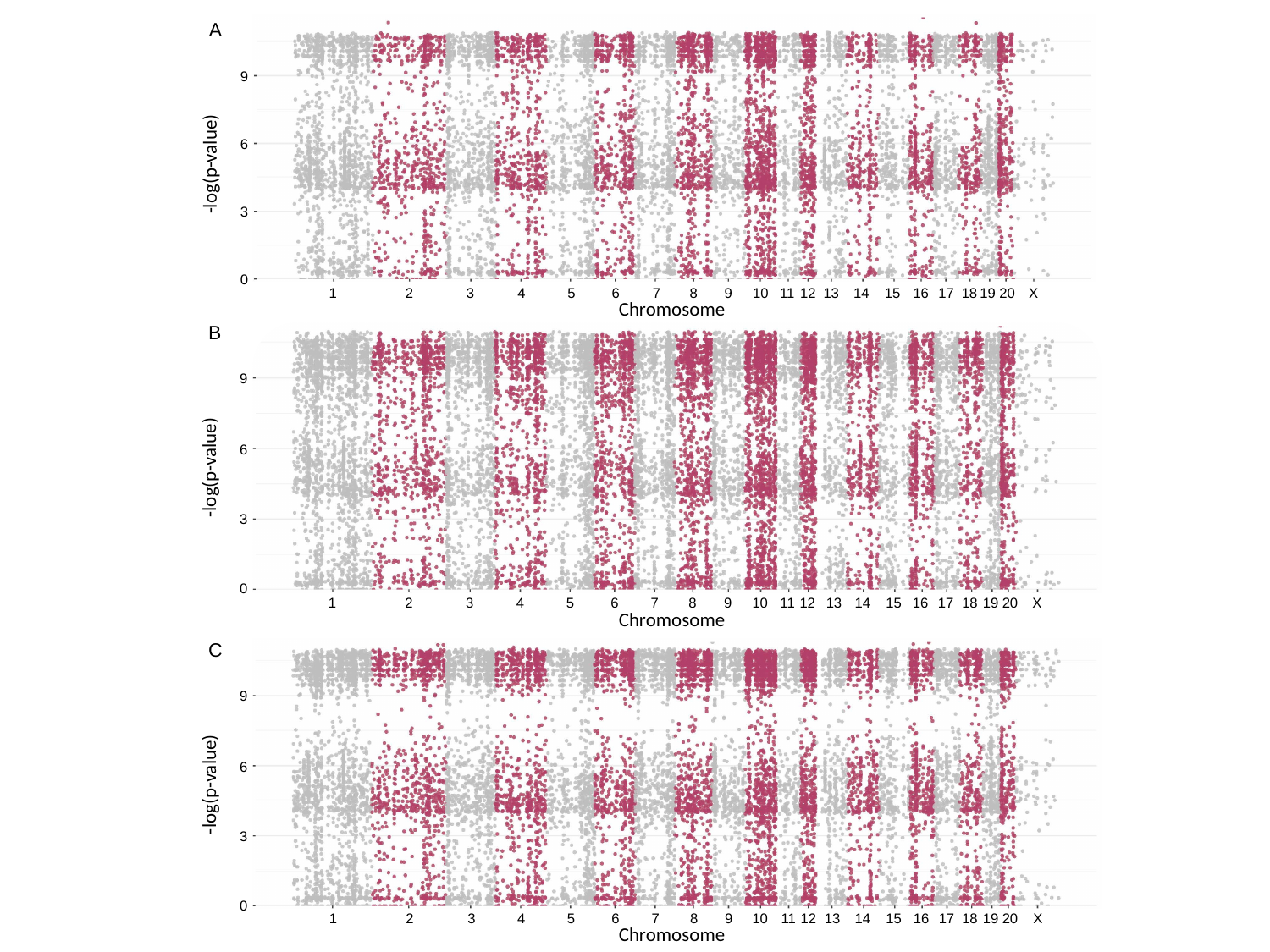

A
9
-log(p-value)
6
3
0
1
2
3
4
5
6
7
8
9
10
11
12
13
14
15
16
17
18
19
20
X
Chromosome
B
9
-log(p-value)
6
3
0
1
2
3
4
5
6
7
8
9
10
11
12
13
14
15
16
17
18
19
20
X
Chromosome
C
9
-log(p-value)
6
3
0
1
2
3
4
5
6
7
8
9
10
11
12
13
14
15
16
17
18
19
20
X
Chromosome

### Slide 3
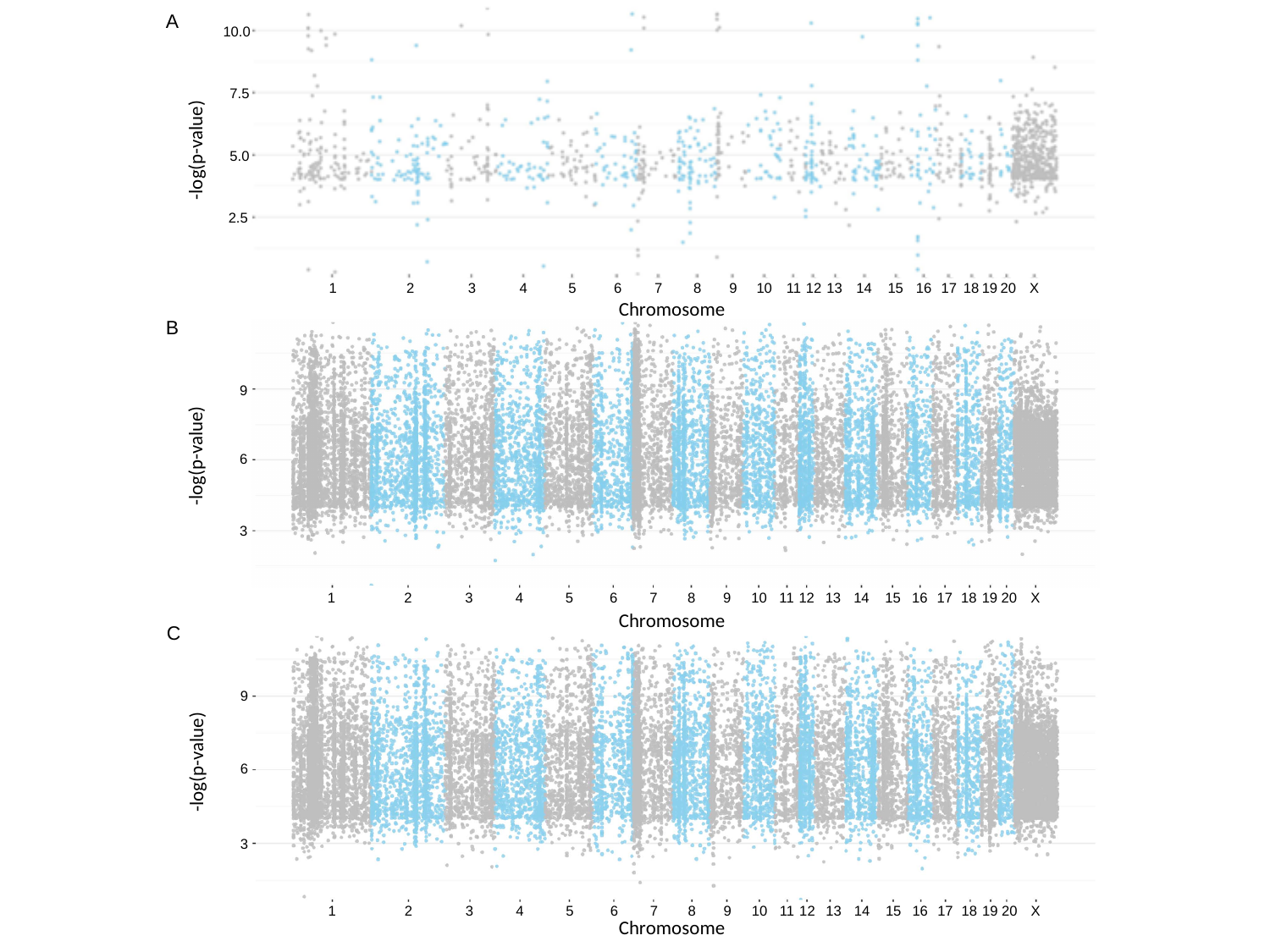

A
7.5
-log(p-value)
5.0
2.5
10.0
1
2
3
4
5
6
7
8
9
10
11
12
13
14
15
16
17
18
19
20
X
Chromosome
B
9
-log(p-value)
6
3
1
2
3
4
5
6
7
8
9
10
11
12
13
14
15
16
17
18
19
20
X
Chromosome
C
9
-log(p-value)
6
3
1
2
3
4
5
6
7
8
9
10
11
12
13
14
15
16
17
18
19
20
X
Chromosome
